# Biophysical Profiling of Red Blood Cells from Thin-film Blood Smears using Deep Learning

**DOI:** 10.1101/2024.04.10.588926

**Authors:** Erik S. Lamoureux, You Cheng, Emel Islamzada, Kerryn Matthews, Simon P. Duffy, Hongshen Ma

## Abstract

Microscopic inspection of thin-film blood smears is widely used to identify red blood cell (RBC) pathologies, including malaria parasitism and hemoglobinopathies, such as sickle cell disease and thalassemia. Emerging research indicates that non-pathologic changes in RBCs can also be detected in images, such as deformability and morphological changes resulting from the storage lesion. In transfusion medicine, cell deformability is a potential biomarker for the quality of donated RBCs. However, a major impediment to the clinical translation of this biomarker is the difficulty associated with performing this measurement. To address this challenge, we developed an approach for biophysical profiling of RBCs based on cell images in thin-film blood smears. We hypothesize that subtle cellular changes are evident in blood smear images, but this information is currently undetectable by human cognition. To test this hypothesis, we developed a deep learning strategy to analyze Giemsa-stained blood smears to assess the subtle morphologies indicative of RBC deformability and storage-based degradation. Specifically, we prepared thin-film blood smears from 27 RBC samples (9 donors evaluated at 3 storage timepoints) and imaged them using high-resolution microscopy. Using this dataset, we trained a convolutional neural network to evaluate image-based morphological features related to cell deformability. The prediction of donor deformability is strongly correlated to the microfluidic scores and can be used to categorize images into specific deformability groups with high accuracy. We also used this model to evaluates differences in RBC morphology resulting from cold storage. Together, our results demonstrate that deep learning models can exceed the limits of human cognition to detect subtle cellular differences in morphology resulting from deformability and cold storage. This result suggests the potential to assess donor blood quality from thin-film blood smears, which can be acquired ubiquitously in clinical workflows.

## Introduction

Thin-film blood smear analysis is a routine diagnostic procedure used to evaluate multiple red blood cell (RBC) pathologies including parasitic infections (e.g. malaria), blood cell disorders (e.g. anemia), and hemoglobinopathies (e.g. sickle cell disease and thalassemia).^1^ Blood smears are simple and cost-effective to prepare; they can be imaged using standard microscopes or slide scanners; and they provide a near-permanent record that can be re-analyzed multiple times without loss of fidelity. These characteristics make blood smear analysis a favourable approach for large-scale clinical studies. Blood smears are currently analyzed by human reviewers to identify morphological features, such as sickle cell and parasitic infections, that can be easily recognized using human cognition. This analysis could then be automated by training AI models on data assessed by expert human reviewers.^2,3^ An emerging possibility in this field, however, is the potential to develop AI models that could detect features from blood smears that cannot be identified by human cognition.

RBC deformability is a compelling physical biomarker of RBC dysfunction, as the loss of RBC deformability is frequently observed in sickle cell disease,^4^ thalassemia,^5^ and infectious diseases like malaria.^6,7^ RBC deformability is also important in transfusion medicine since the loss of RBC deformability is a consequence of RBC senescence^8^ and the RBC storage lesion.^9–12^ Transfusing less deformable RBCs is less effective because the cells fail to transit the microvasculature and are instead removed from circulation in the spleen,^13,14^ resulting in a shorter circulation time and the need for more frequent transfusion.^15^ Interestingly, RBCs from some donors seem to resist RBC damage during storage, suggesting that these donor RBCs could provide longer-circulating transfusion units.^9,10^ These long-circulating RBC units could be reserved for chronic transfusion recipients to enhance the interval between transfusions, reducing the frequency of transfusion as well as the risk for transfusion-associated adverse events.^9,16–19^

In contrast to the standard blood smear analysis, measuring RBC deformability requires specialized equipment and expert personnel. Available methods include flow-based methods, such as ektacytometry;^20,21^ micro-mechanical deformation methods, such as micropipette aspiration,^22^ atomic force microscopy,^23^ magnetic twisting cytometry,^24–26^ electrodeformation,^27–29^ and optical tweezers;^30,31^ as well as microfluidic methods, such as deformation in microchannels,^32,33^ electrical impedance sensing,^34^ deformation through capillaries,^35^ constrictions,^36–42^ tapered constrictions,^43–45^ and sorting by microfluidic ratchet.^46–48^ Further, imaging approaches have been integrated with microfluidics, developing into the emerging field of optofluidics.^49^ Alongside optofluidics, another imaging approach to evaluate RBC deformability is quantitative phase imaging (QPI).^50–56^ These imaging approaches vastly improve the throughput for RBC deformability assessment.^49,52^ However, optofluidics and QPI suffer similar drawbacks as other RBC deformability assessment methods, including the need for expensive equipment, highly skilled personnel, and precise experimental procedures. All of these measurement methods are currently prohibitively laborious in routine clinical analysis.^57^

Here, we investigate the potential to assess RBC deformability from microscopy imaging of thin-film blood smears to provide a potential path for routine clinical assessment of this physical biomarker. Our method involves measuring RBC deformability using the microfluidic ratchet device while simultaneously imaging a blood smear from the same sample to train a convolutional neural network (CNN) to infer RBC deformability. We demonstrate that deformability can be evaluated using CNN models to perform both regression (*r*=0.708, *p* < 10^-6^) and classification (79%). Furthermore, a CNN model used to detect weekly storage duration achieved 89 ± 8% testing accuracy for individual donors and 78% for pooled donor datasets. Importantly, we observed that this CNN model can be generalized to evaluate previously unseen samples from new donors. These results demonstrate the potential to assess blood smears to screen for donors that can provide resilient RBCs that retain their deformability during storage and are likely to circulate longer in recipients. These long-circulating RBCs could then be preferentially used by chronic transfusion recipients to extend the interval between transfusions and thereby reduce the occurrence of transfusion-related morbidities, such as iron-overload.

## Results

### Approach

Our experimental approach involves acquiring RBC samples from multiple donors and storing them in standard test tubes at 4℃ for 0, 1, and 2 weeks to generate samples with a range of deformabilities by accelerating their aging (**Fig. 1A**).^9,58^ The microfluidic ratchet cell sorting mechanism is used to precisely measure RBC deformability on each sample based on their ability to deform through a matrix of constrictions to separate cells into distinct deformability outlets (**Fig. 1B**).^36,46,47^ The sorted cell distribution is then used to obtain a rigidity score (RS) for each sample (**Fig. 1C**). In parallel, a thin film blood smear is prepared from each RBC sample and imaged using brightfield microscopy at 60× (**Fig. 1D-E**). The microscopy images are segmented into a standard size in order to create 10,000 randomly selected non-overlapping images to train a supervised convolutional neural network (**Fig. 1G**). The ground-truth labels are obtained from the deformability measurement results obtained for each sample. This work improves upon our previous work, which analyzed live RBC images. Here, we analyzed Giemsa-stained thin-film blood smears, which provide greater morphological data with less sensitivity to sample preparation and environmental conditions. Specifically, we generated two separately labelled datasets for CNN evaluation of RBC morphology based on (1) deformability and (2) storage time.

**Fig. 1.**
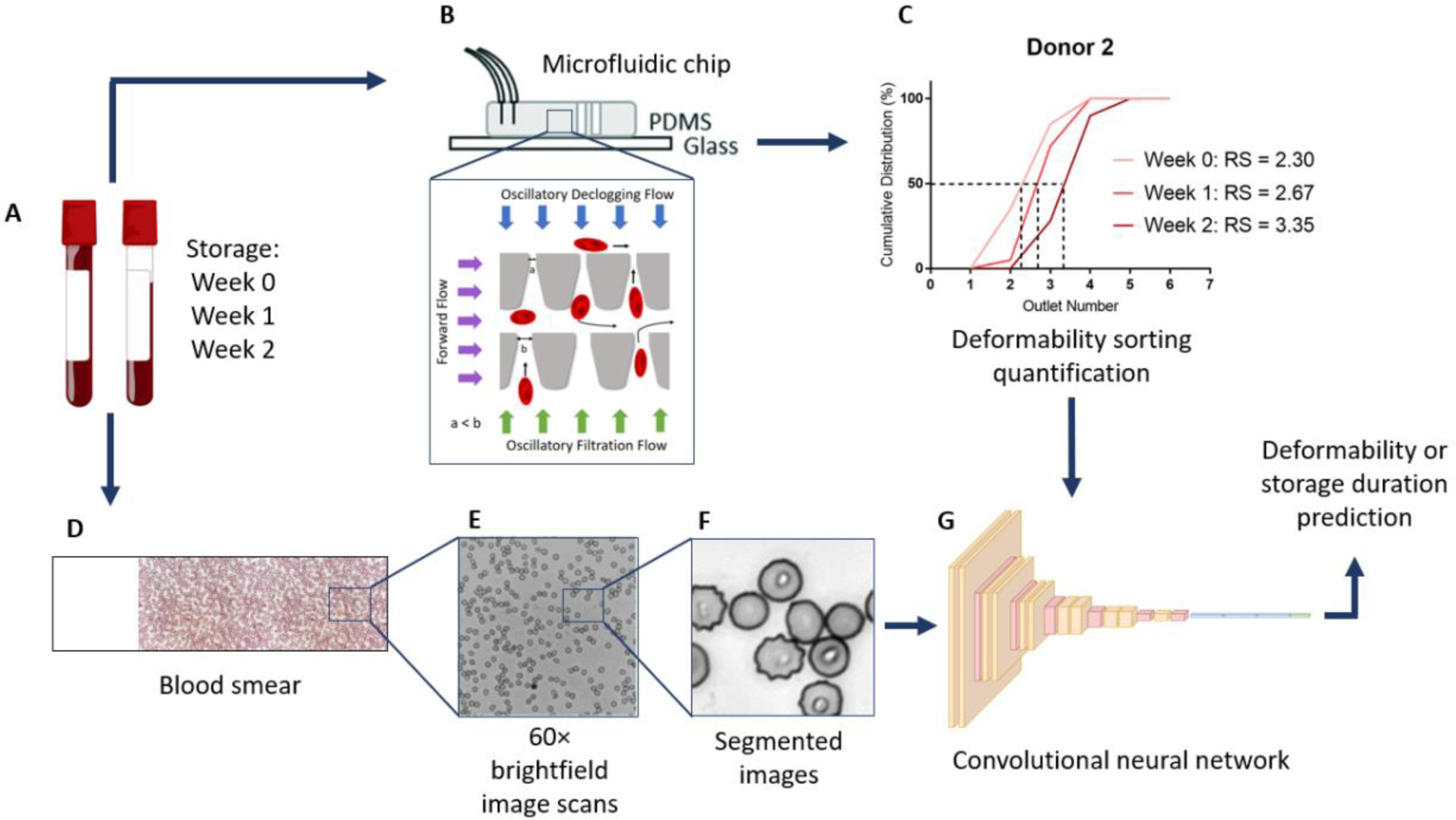
Overall experimental approach. (A) Processed blood samples undergo cold storage in tubes for an accelerated damage storage model. (B) A microfluidic ratchet sorting device assesses the donor’s RBC deformability profile and degradation during ageing. (C) A rigidity score (RS) is assigned to each RBC population. (D) Concurrent to the microfluidic evaluation, stored blood is smeared onto glass slides, fixated with methanol, and stained with Giemsa. (E) The smears are imaged with an optical microscope in brightfield at 60× magnification. (F) The captured images are segmented into 256 × 256 pixel images containing multiple cells. (G) The segmented images are processed into training and testing sets for deformability and storage duration prediction using a convolutional neural network.

### Measurement of RBC deformability

To obtain the training labels for image-based prediction of RBC deformability, we used a microfluidic ratchet device to measure RBC deformability. We collected RBCs from nine healthy donors and stored these cells in standard polypropylene tubes to promote the accelerated loss of cell deformability.^9^ We assessed these cells at weeks 0, 1, and 2 to establish the RS for each RBC sample. RBC deformability was measured using the previously established microfluidic ratchet device.^9,59^ Briefly, the microfluidic ratchet device uses oscillatory flow of RBCs through a matrix of progressively narrowing tapered constrictions, depositing cells into twelve deformability outlets (Fig. 2A). Based on the microfluidic design, fluid pressures, and cell mechanics, the majority of RBCs are sorted into outlets 2-5 at a throughput of ∼600 cells per minute. To characterize the deformability of the RBC sample, the sorted distributions are plotted as a linearly interpolated cumulative distribution. A rigidity score (RS) can then be obtained from the outlet value at the 50% crossover point (Fig 2B).^9,59^

**Fig. 2.**
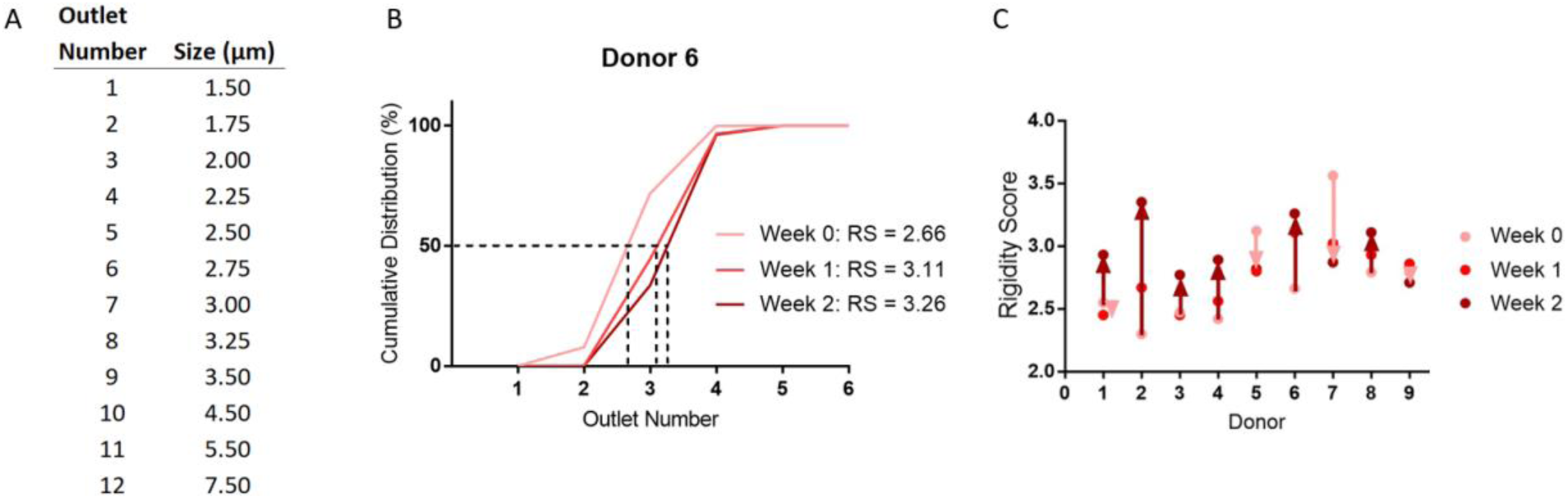
Donor rigidity scores (RS) assessed by the microfluidic ratchet sorting device. (A) Microfluidic funnel constriction size for each outlet number. (B) Example deformability sorting results over the storage period for Donor 6. The RS is defined as the outlet number at the 50% crossover frequency of the cumulative distribution and correlates with greater cell rigidity. (C) Donor rigidity score aging profiles. We observe a general trend of cell rigidification during storage (6 donors), of which four exhibited major RBC rigidification and two with minor rigidification (dark red arrows). There were also two donors displaying minor RBC softening and one with substantial softening during storage (light red arrows).

Initially, we conducted cell sorting using chemically degraded RBCs to validate the microfluidic ratchet for detecting deformability degradation. We used glutaraldehyde (GTA) to chemically degrade an RBC sample over five cases: untreated control RBCs and treated cells with GTA concentrations of 0.010, 0.0125, 0.015, and 0.025%. We find a predictable decrease in cell deformability with increasing GTA concentration (Fig. S2). This provides us with evidence that the microfluidic device can detect sensitive changes to RBC deformability over the course of storage.

Ultimately, we observed a general trend of increasing RS for donor RBCs tested over two weeks of storage in test tubes. This result is consistent with the loss of RBC deformability that occurs during accelerated aging from the RBC storage lesion (Fig. 2C and Fig. S3). Consistent with our previous studies, we observed donor-dependent variability in fresh RBC deformability and differences in RBC rigidification in response to cold storage.^9^ These results provide samples with a range of RBC deformability as the training data set for image analysis of blood smears.

### Optical microscopy and image processing

To generate an image dataset for deep learning analysis, we generated collections of unbiased image segments from all donors at each experimental time point. For each RBC sample, we created a thin-film smear that was fixed with methanol and stained with Giemsa to enhance contrast and highlight morphological variations. Using a 60× magnification brightfield contrast objective, we captured high-quality, consistent images of these fixed and stained smears, yielding a total of 432 large images, each with a resolution of 2424 × 2424 pixels (Fig. 3A). We segmented each large image into 81 non-overlapping sections of 256 × 256 pixels, resulting in 34,992 segmented images per class (Fig. 3B). We used the pixel intensity of cell-free images as a threshold to exclude and remove empty segmented images. The selected images contained considerable variations in the number, position, and density of RBCs. Images with substantial cell overlap or occlusion were filtered and removed. This image screening procedure provided the model with an unbiased dataset containing a variety of image examples, enabling the model’s learning capability for evaluating unstructured blood smear images.

**Fig. 3.**
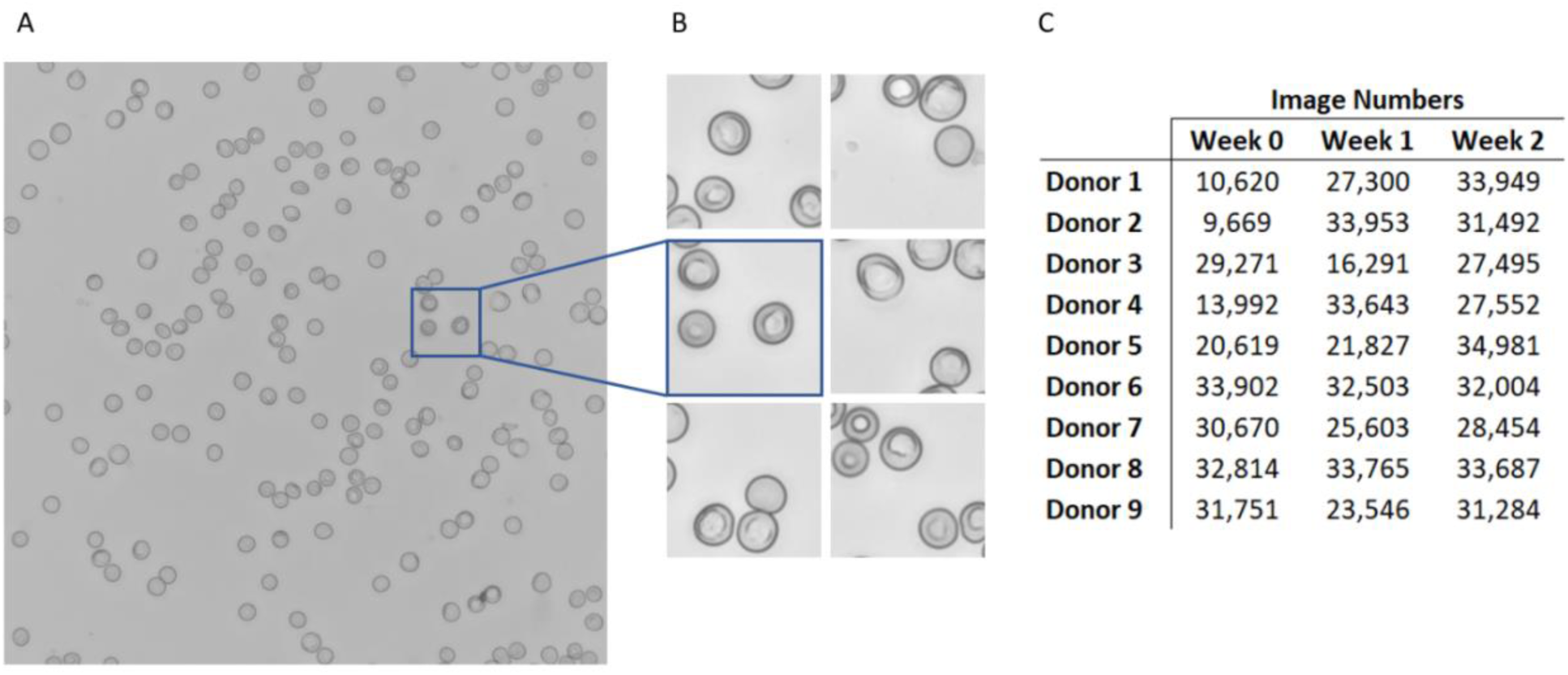
Image acquisition. (A) Captured blood smear image scan of 2424 × 2424 pixels and (B) segmented 256 × 256 pixel images from Donor 3, after storage for week 1. (C) Number of images acquired from each donor at different storage time points after filtering. These images were used to create training, validation, and testing datasets.

Following the segmentation and filtering of images, we generated image collections to serve as training and testing sets for a convolutional neural network (CNN) model. We partitioned the filtered images into an 80:20 ratio for the training and testing sets, to produce a training set of 10,000 images and a testing set of 2,000 images for each smear. We randomly selected images from each set and rotated them by a multiple of 90°, reducing potential biases related to image orientation and lighting artefacts. Since our image processing workflow generated images in excess of the cells needed to generate this 12,000 image dataset (Fig. 3C), we randomly sub-sampled the acquired data.

### Network design, training, and validation

Using the datasets generated from imaging blood smears, we designed and tested a CNN to conduct image feature extraction and classification (Fig 1G). Using the PyTorch library, we generated a neural network model comprised of a series of five convolutional layers and five max pooling layers, which work together for feature extraction. The network begins with an initial convolutional layer with a kernel size of 7 × 7. This is followed by a layer with a 5 × 5 kernel size, and then three layers with 3 × 3 kernel size. All convolutional layers use a stride of 1 and “same” padding. This type of padding adds zero-valued pixels to the image borders, ensuring that the entire image is covered by the filter. Each convolutional layer is followed by rectified linear unit (ReLU) activation, and batch normalization. By using five convolutional layers, the model captures low-level features in the early layers (i.e., edges and corners) and high-level features in the later layers (i.e., shapes and patterns). Furthermore, by padding the image edges we can ensure that the RBCs randomly distributed throughout the entire image segment, including the edges, are incorporated into the model.

After feature extraction, the network transitions to the classification section, which is comprised of three fully connected layers. Each of these layers is followed by ReLU activation, batch normalization, and 20% dropout. The output layer, dedicated to classification, employs a SoftMax error function for backpropagation during training. For optimization, the network utilizes cross-entropy loss and stochastic gradient descent. Finally, we trained the CNN using 10,000 multi-cell images (256 × 256 pixels) per class from each donor that were collected at weeks 0, 1, and 2 of cold storage. Influenced by the AlexNet architecture and as described in our previous work,^59,60^ this CNN model is used to associate RBC images with their biophysical and morphological characteristics.

### Classification based on cell deformability

Our previous work demonstrated a two-class classification of RBCs under precise live-cell imaging conditions.^59^ Here, we adapted the CNN analysis to predict RBC deformability from blood smear images. We classified RBCs into four classes, defined by RS values: RS<2.50, RS 2.50-2.74, RS 2.75-2.99, or RS>2.99 (Table 1). In total, 50,000 images per class were used for training and 10,000 images per class were used for testing.

**Table 1.** Binned deformability categories for each donor and storage time.

| Donor | RS < 2.50 | RS 2.50-2.74 | RS 2.75-2.99 | RS > 2.99 |
| --- | --- | --- | --- | --- |
| 1 | Week 1 | Week 0 | Week 2 | - |
| 2 | Week 0 | Week 1 | - | Week 2 |
| 3 | Week 0, 1 | - | Week 2 | - |
| 4 | Week 0 | Week 1 | Week 2 | - |
| 5 | - | - | Week 1, 2 | Week 0 |
| 6 | - | Week 0 | - | Week 1, 2 |
| 7 | - | - | Week 2 | Week 0, 1 |
| 8 | - | - | Week 0, 1 | Week 2 |
| 9 | - | Week 2 | Week 0, 1 | - |
| Count | 5 | 5 | 10 | 7 |

**Table 2.** Testing metrics for donor deformability classification.

| Model | Testing Accuracy | Precision | Recall | F1-Score |
| --- | --- | --- | --- | --- |
| Deformability classification | 0.79 | 0.798 | 0.786 | 0.779 |

This analysis achieved 79% testing accuracy. Misclassified cells were largely at the interface between neighbouring classes. This outcome suggests that the model learns common deformability features between donors and observes similarities in images near the class boundaries (Fig. 4A). The saliency maps obtained for this analysis revealed that the features used for classification were located within the cell (Fig. 4B). Direct imaging of RBCs from each of the deformability classes showed a progressively increasing frequency of more rigid echinocyte RBCs (Fig. 4C). The increased frequency of echinocytes during storage could contribute to the increased RS since these cells have a decreased surface area to volume ratio compared to discocytes.^61,62^

**Fig. 4.**
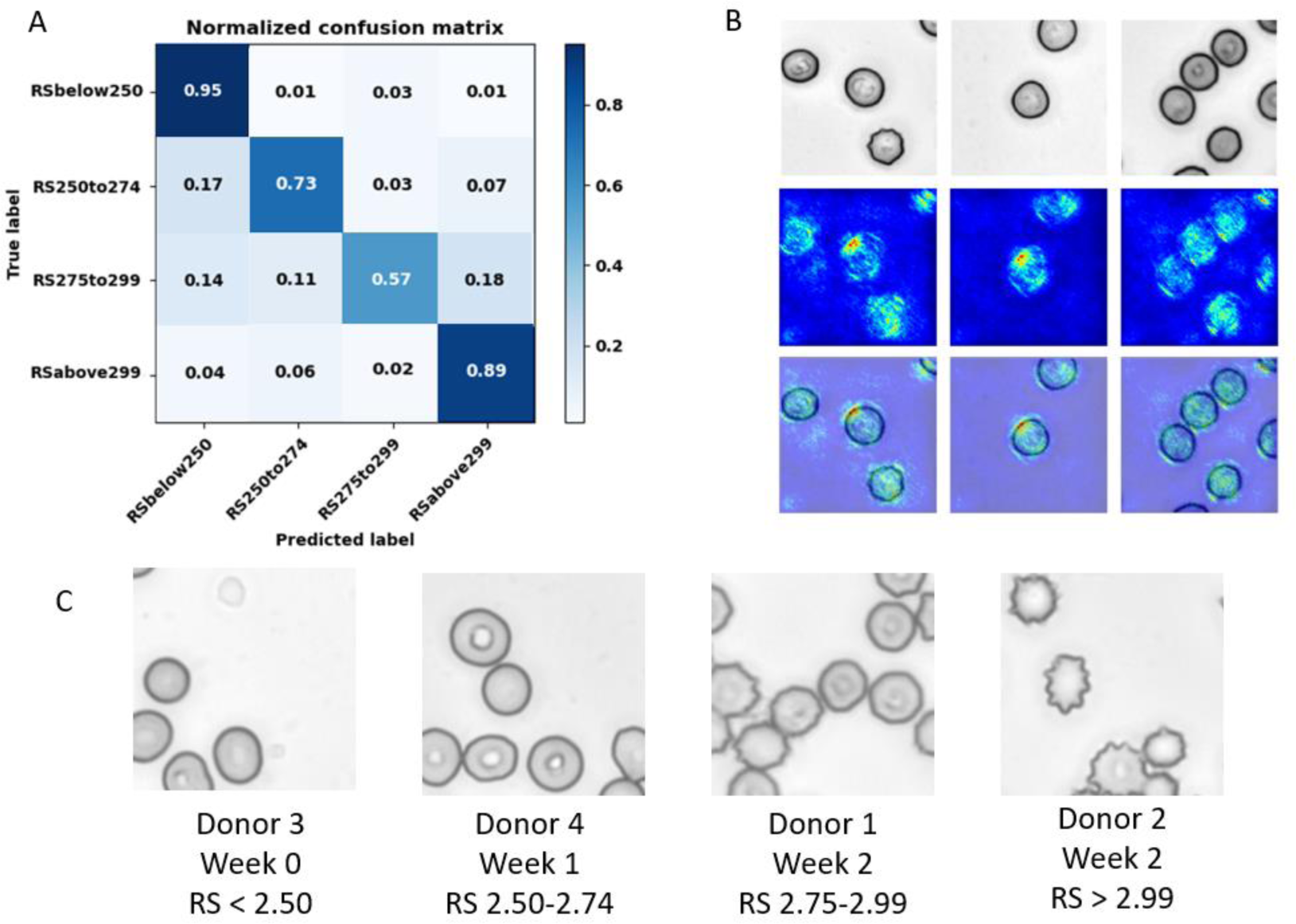
Collective donor deformability classification. (A) Normalized confusion matrix of RBC deformability classification. This model was trained using the training datasets from each donor and tested on the held-out testing datasets. We achieved 79% testing accuracy in this analysis. (B) Top: example brightfield input images. Middle: saliency maps. Bottom: saliency maps overlayed on the input images. (C) Example images from each deformability class.

### Cell deformability regression analysis

We performed linear regression analysis to associate the true RS (measured by microfluidic ratchet) with the RS predicted by the CNN. Image datasets were associated with 27 labels, each representing image collections from nine donors at weeks 0, 1, and 2. To adapt CNN classification for regression analysis, we used a custom data loader to assign continuous regression labels, modified the final layer to output a single channel representing the predicted RS, and optimized the network for Mean Squared Error (MSE) loss. After tuning the hyperparameters, the model exhibited robust performance on the testing dataset, where the Mean Absolute Error (MAE) shows that the predicted RS deviates an average of 0.164 from the measured value (Table 3). This degree of deviation is consistent with the RS generated by the microfluidic device, which displays a variation of ± 0.17 for the same donor during subsequent runs on the same day.^9^ The Root Mean Squared Error (RMSE) of 0.212 is higher than the MAE, as it emphasizes predictions that substantially deviate from the target. However, the RMSE is not considerably greater than the MAE, suggesting that this deviation was a consequence of data outliers. Together, over the range of RS values (2.30 to 3.56), these errors correspond to a percentage deviation of 13.5% for microfluidic variation, 13.0% for MAE, and 16.8% for RMSE. Furthermore, there was a strong correlation between the predicted and true microfluidic RS values (*r* = 0.708) and the pixels contributing to the predictions were predominantly within the cell (Fig. 5). This result indicates that the CNN model predicts a donor’s RS value at a level of accuracy comparable to the microfluidic ratchet measurement.

**Fig. 5.**
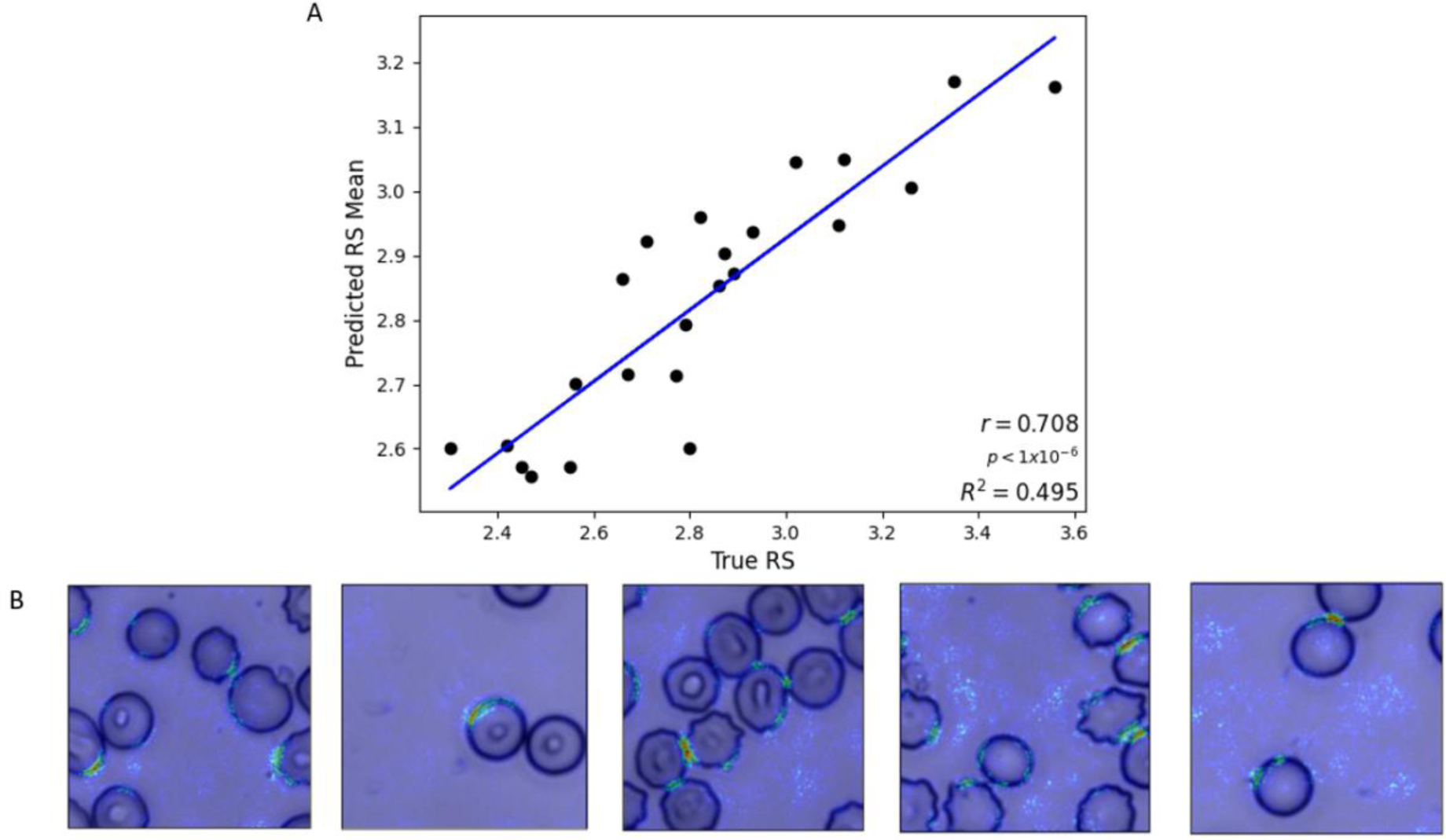
Donor deformability regression results. (A) The mean predicted rigidity score is plotted vs the true rigidity score. (B) Testing saliency maps using warm colouration to depict highly weighted pixels for prediction, demonstrating their concentration around the cells.

**Table 3.**
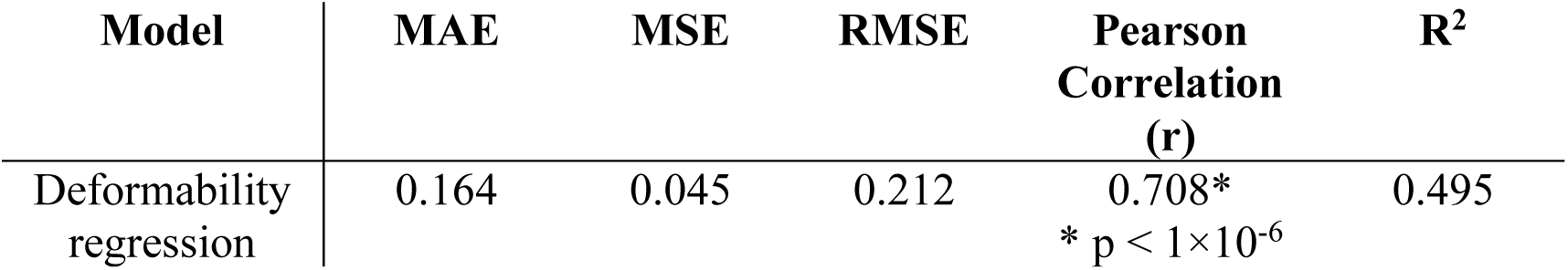
Testing performance metrics for RS deformability prediction using a regression model. All donors are represented in the training and test sets.

| Model | MAE | MSE | RMSE | Pearson Correlation (r) | R<sup>2</sup> |
| --- | --- | --- | --- | --- | --- |
| Deformability regression | 0.164 | 0.045 | 0.212 | 0.708* | 0.495 |
| * $p < 1 \times 10^{-6}$ | | | | | |

### Individual donor storage duration classification

We applied our CNN model to analyze datasets from individual donors to predict storage duration based on RBC images over two weeks of storage (Fig. 6A). We observed evidence of the storage effect within the first two weeks, including the emergence of increased frequency of echinocytes recognizable due to their evenly spaced thorny cell membrane projections. We trained a CNN classification model and established validation accuracies averaged over five folds, using 10,000 images from each donor on week 0, 1, and 2. Subsequently, we performed testing with 2,000 images per class that are collected from the same donor and reserved for testing. We observed comparable levels of accuracy between the training, validation, and testing stages for each donor, obtaining an overall average testing accuracy of 89 ± 8% across the donors (Fig 6B). The saliency maps confirmed that the pixels contributing to these predictions are concentrated within the cells (Fig. 6C). Additional testing metrics including precision, recall, and F1-score, are available in the Supplementary Materials (Table S1).

**Fig. 6.**
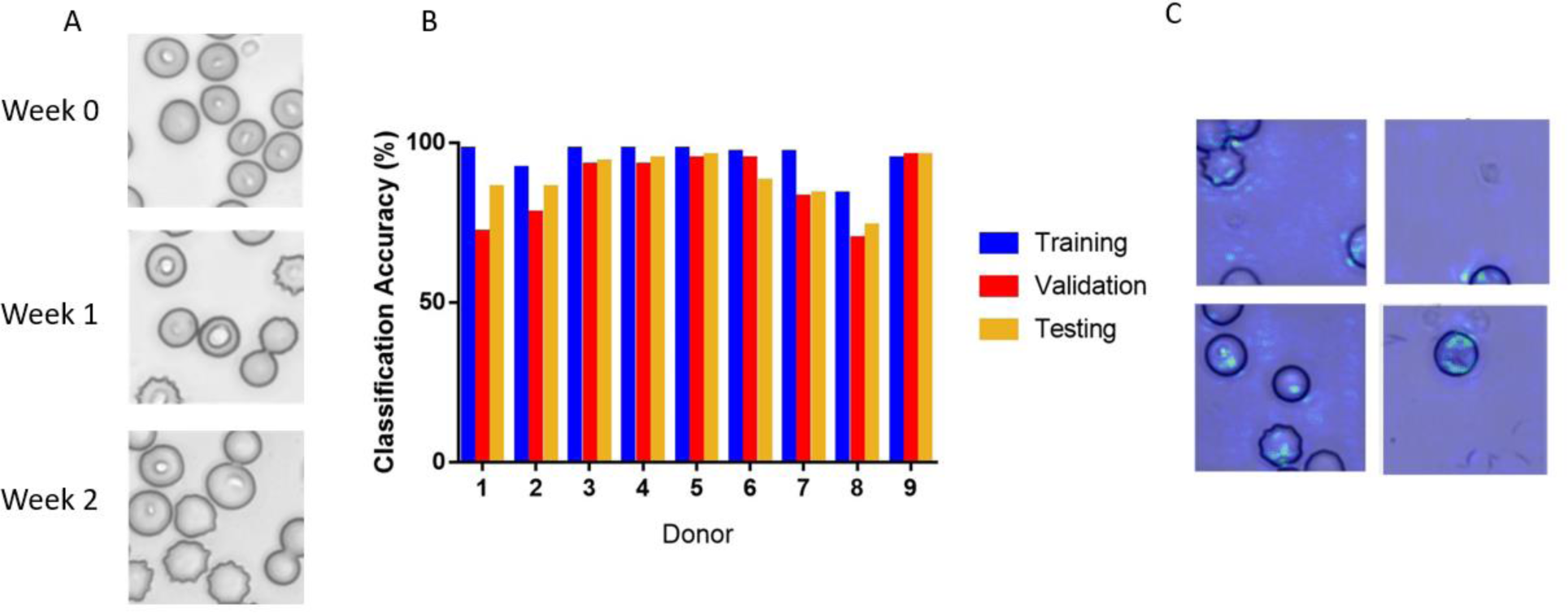
Individual donor storage duration classification. (A) Example images from Donor 2 for storage weeks 0, 1, and 2. Note the morphological changes in weeks 1 and 2 compared to week 0. (B) Training, validation, and testing accuracies of storage duration prediction for individual donors. The aggregate average of testing accuracy was 89 ± 8%. (C) Example saliency maps from Donors 2, 4, and 5.

### Collective donor storage duration classification

Following the individual analyses, we combined RBC images from multiple donors to develop a generalizable model to predict the storage time of RBCs from donors that were not included during CNN training. First, we combined all donor RBC images into a single dataset of 270,000 images (10,000 per class per donor). We tested the CNN using 2,000 reserved images per class per donor, for 54,000 testing images total. From the testing dataset, we generated a normalized confusion matrix (Fig. 7A) and achieved 87% training accuracy and 78% testing accuracy (Table 4). Since all donors were used in training and testing here, this accuracy represents the upper-bound for subsequent testing with donor datasets not used in training. Saliency maps, illustrating the clustering of activated pixels around the cells (Fig. 7B), support the idea that our model learns cell-based morphological features. Next, we repeated this process but generated a training dataset from all donors save one. The excluded donor dataset was used to test whether the CNN could predict storage duration of RBCs from donors upon which they were not trained. This analysis achieved an aggregate testing accuracy of 61 ± 7% (Table 4), which is still a notable result considering the challenges associated with developing generalizable predictive models.

**Fig. 7.**
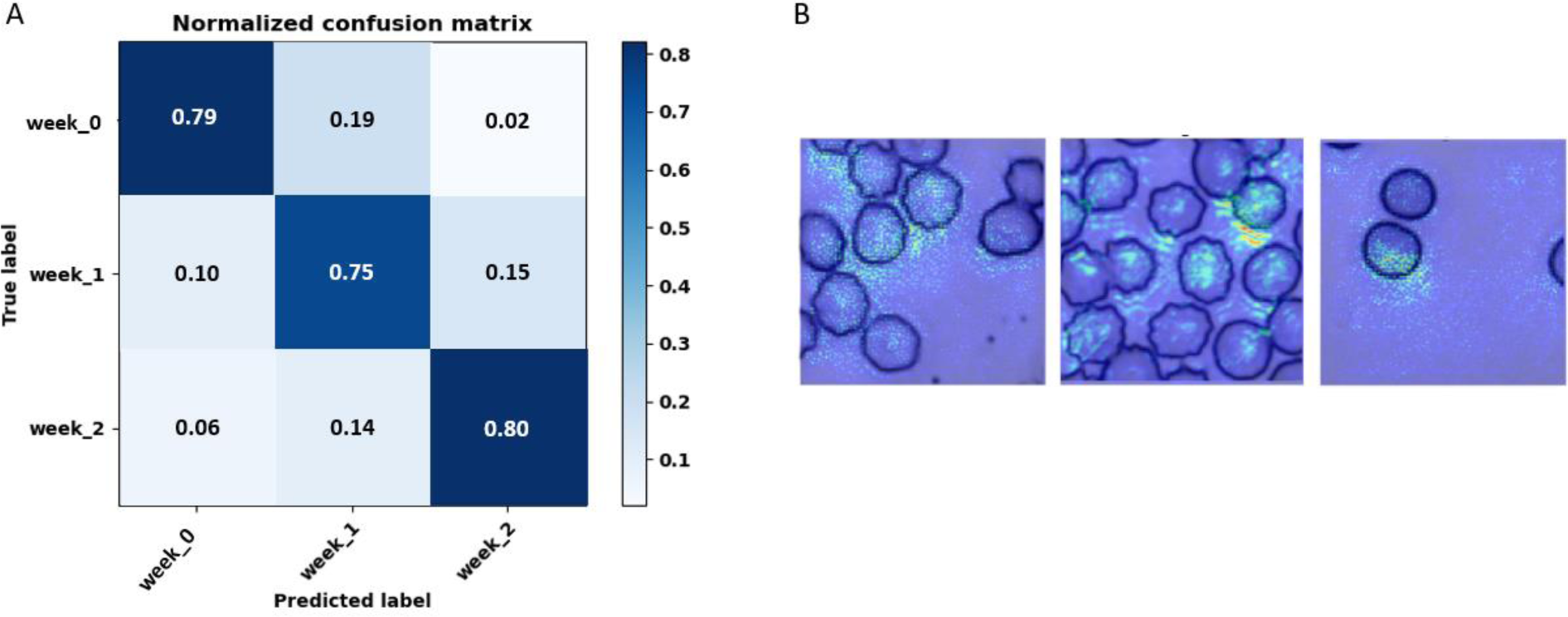
Collective donor storage classification. (A) Normalized confusion matrix of storage duration prediction on the testing set containing all donors. The training accuracy was 87% and the testing accuracy was 78%. (B) Saliency maps indicating the pixels that contributed the most to classification.

**Table 4.** Testing performance metrics for storage time prediction using a classification model. For the first case, all donors are represented in the training set. In subsequent cases, the donor selected for testing is held-out from the training set.

Red Cell Quality from Blood Smears
| Testing Donors | Testing Accuracy | Precision | Recall | F1-Score |
| --- | --- | --- | --- | --- |
| All donors | 0.78 | 0.869 | 0.868 | 0.868 |
| Donor 1 | 0.75 | 0.835 | 0.858 | 0.833 |
| Donor 2 | 0.60 | 0.629 | 0.640 | 0.626 |
| Donor 3 | 0.53 | 0.557 | 0.570 | 0.561 |
| Donor 4 | 0.55 | 0.499 | 0.545 | 0.499 |
| Donor 5 | 0.66 | 0.628 | 0.657 | 0.639 |
| Donor 6 | 0.57 | 0.563 | 0.569 | 0.555 |
| Donor 7 | 0.70 | 0.667 | 0.674 | 0.669 |
| Donor 8 | 0.58 | 0.685 | 0.676 | 0.678 |
| Donor 9 | 0.58 | 0.557 | 0.538 | 0.544 |
| Average | 0.61 | 0.624 | 0.636 | 0.623 |
| Standard deviation | 0.07 | 0.099 | 0.100 | 0.100 |

## Discussion

This study investigated machine learning analysis of Giemsa-stained thin-film blood smears as an alternative to direct biophysical measurement of RBC deformability. The advantage of blood smear analysis over direct biophysical measurement is that blood smears are part of the standard clinical laboratory workflow, which enables measurement of RBC deformability without the need for specialized equipment and expert personnel. Analysis of RBC images using a CNN model enabled classification of RBCs into four categories of cell rigidity with 79% classification testing accuracy. Using a regression model, we were also able to predict specific RS values for each test image with a mean absolute error of 13.0%, which is comparable to the mean absolute error of 13.5% for direct measurement using the microfluidic ratchet device.^9^

Our previous work involved imaging RBCs extracted from the outlets of a microfluidic ratchet device, which sorts cells based on deformability, and used a CNN model to classify cells in to into two deformability groups.^59^ The work presented here entirely removes the need to separately analyze sorted RBCs by inferring deformability from Giemsa-stained slides. The use of microscopy slides is important because fast and low-cost automated imaging workflow solutions are already commercially available.^63^ We enhanced the analytic approach to classify cells into four deformability groups and subsequently performed continuous regression prediction in order to infer a precise rigidity score for the RBC population. The enhancements to the methods enabled the CNN classification model to predict storage duration (weeks 0, 1 and 2) using individual donor datasets and combined donor datasets, with high accuracy for individual (89%) and collective (78%) storage duration prediction. Furthermore, when a donor was held-out from training and was reserved for testing to establish that the model was generalizable for storage duration prediction, this produced greater accuracy (53-75%) than random chance (33%). Future work can incorporate larger datasets to improve the generalizability of this method to assess RBC deformability.

This image analysis approach used here provide a significant advance over direct biophysical measurement of RBC deformability. Microfluidic assessment of RBC deformability has become a preferred method because of high throughput over conventional approaches, such as micropipette aspiration,^64–66^ optical tweezers,^67–69^ and atomic force microscopy.^23,70^ However, these methods are limited by the need for specialized instruments and expert personnel to assess the deformations of RBCs, which present a significant barrier for clinical analysis of this biophysical parameter.^57,71,72^ The advantages of imaging approaches, including quantitative phase imaging (QPI)^50–56^ and optofluidics,^49^ include greatly increasing the throughput for RBC deformability assessment.^49,52^ However, QPI requires existing laboratory optical microscopes to be converted into quantitative phase microscopes, limiting its rapid deployment to clinical settings.^51^ In addition, QPI could benefit from improvements in high-throughput measurements and data analysis and management systems,^51^ which could be enabled using machine learning techniques.^56^ Regarding opto-microfluidics, these approaches are increasingly implemented alongside deep learning technologies,^49^ but are primarily limited by the need for developing, maintaining, and validating microfluidic devices which require substantial investments in equipment and personnel. In this work, through both high throughput imaging and machine learning, we achieve an imaging throughput of ∼4,000 cells per minute using microscopy and Giemsa-staining protocols that are ubiquitous in research and clinical laboratories. Taken together, RBC deformability assessment using deep learning of thin-film blood smear images is accessible, efficient, less complex, and standardizable.

Other deep learning methods have previously characterized RBC properties. Morphological variations have been used to assess pathology, such as malaria,^2,73–84^ sickle cell disease,^85–90^ thalassemia,^5,91,92^ pyruvate kinase disease,^93^ and COVID-19.^94^ Similarly, morphologic changes have been assessed during the storage lesion, where blood quality deteriorates with cold storage.^50,95^ Recently, our group and others have used imaging and machine learning to infer biophysical properties, such as RBC elasticity,^96^ deformability,^59^ and plasticity,^93^ in the hopes of developing rapid RBC screening to assess the quality of blood units for transfusion evaluation.^71^ Our extension of this strategy to use Giemsa-stained blood smears is an important advance as it synergizes deep learning methods with existing clinical blood quality assessment protocols.

Together, we present RBC imaging and machine learning analysis of standard Giemsa-stained thin blood smears as a rapid, low-cost, and clinically accessible method for measurement of blood quality. This method uses cell morphology to predict the biophysical characteristics of RBCs, such as cell deformability, which are not easily accessible through direct measurement. These methods could be incorporated into existing blood processing workflows to assess the quality of blood as well as to predict the impact of cold storage. By identifying high-quality RBC units, they could be reserved for the vulnerable and chronic transfusion recipients, to improve outcomes of blood transfusions for these individuals while more effectively using the available blood supply.

## Methods

### RBC sample collection, preparation, and storage

This study was approved by the University of British Columbia Clinical Research Ethics Board (UBC REB# H19-01121) and the Canadian Blood Services Research Ethics Board (CBS REB# 2019-029). After informed consent, blood samples were collected in Ethylenediaminetetraacetic acid (EDTA) tubes from each participant. The donors were between the ages of 18-70 and self-identified as healthy.

Whole blood samples were separated by centrifuging at 3180 *g* for 8 minutes. Blood supernatant was removed and stored for further processing. The packed RBC pellet was then centrifuged at 677 *g* for 30 minutes with no brakes to decant the plasma. Each 1 mL of packed

RBCs was mixed with 0.80 mL of saline-adenine-glucose-mannitol storage solution (SAGM) and 0.56 mL of plasma. Aliquots were collected for microfluidic deformability sorting and blood smear within 48h following each storage timepoint: week 0 (day 0), week 1 (day 7) and week 2 (day 14). For microfluidic deformability sorting, the aliquot was resuspended and washed three times in a five-fold dilution of Hanks balanced salt solution (HBSS, Gibco) with 0.2% Pluronic solution (F127, MilliporeSigma). The washed RBC pellet was suspended in HBSS at <1% hematocrit for infusion into the microfluidic device.

For the thin-film blood smear, 10 µL of the storage solution was placed on a slide and smeared across with another slide angled at 45° from the surface. After drying, the smear was dipped in methanol to fix the cells to the surface and to maintain their morphology. After the methanol dried, the smear was stained with Giemsa stain in an Coplin jar for 20 minutes. After staining, the smear was washed three times in deionized water.

### Microfluidic ratchet device manufacture

The manufacture of the microfluidic devices has been described previously.^97,98^ We created a master mold for the device using photolithographic microfabrication. This was used to create a secondary master polyurethan mold fabricated from Smooth-Cast urethan resin (Smooth-Cast ONYX SLOW, Smooth-On), described in more detail here.^99^ We produced single-use microfluidic devices from the secondary master mold using PDMS silicone (Sylgard-184, Ellsworth Adhesives) mixed at a 10:1 ratio with the PDMS curing agent (Sylgard-184, Ellsworth Adhesives). After the PDMS was mixed and poured into the secondary masters, they were cured for two hours at 65°C. After curing, the PDMS molded devices were removed from the molds and were manually punched with 0.5- and 3.0-mm hold punches (Technical Innovations). Then, a thin PDMS silicone (RTV 615, Momentive Performance Materials LLC) layer was mixed at 10:1 ratio with the RTV 615 curing agent and spun into a thin layer on a 100 mm diameter silicone wafer (University Wafer) at 1500 rpm for 1 minute. The spun layer was then cured for two hours at 65°C. The Sylgard-184 PDMS microstructure mold was bonded to the RTV 615 thin PDMS layer using air plasma (Model PDC-001, Harrick Plasma). This bonding with the thin PDMS layer seals the device’s microstructures. Finally, the composite sealed microstructure mold was then bonded to a 75 × 50 mm glass slide (Corning) using air plasma.

### Microfluidic device operation

The microfluidic ratchet sorting device uses microscale funnel constrictions to measure cell deformability. The mechanism and operation of this device has been described and validated previously.^10,59^ The device is operated using four pressurized fluidic inputs: sample inlet flow, horizontal cross-flow, oscillating upward filtration, and downward declogging flows. The sample flows into the device *via* the sample inlet flow and proceeds to the matrix sorting region where the cells are sorted using a ratcheting effect. The sorting matrix of micropores has openings starting at 7.5 μm for the most rigid outlet (12), decreasing to 1.5 μm for the most deformable outlet (1) (see Fig. 2A for the constriction width of each outlet). The resolution of these pores is limited by the microfabrication process as the resolution of our mask is limited to 250 nm and our photolithography wavelength is limited to 340 nm. Therefore, the smallest possible feature resolution possible with our fabrication process is ∼250 nm.

The device is prepared by buffering HBSS with 0.2% Pluronic-F127 solution through the horizontal crossflow inlet at high pressure (300 mbar) for 15 minutes. The RBC sample at <1% hematocrit suspended in HBSS with 0.2% Pluronic-F127 is input to the sample inlet of the microfluidic device at 40-45 mbar. The horizontal crossflow operates at 55-50 mbar, while the oscillating up and down pressures operate at 175 and 162 mbar, respectively.

### Microfluidic sorting validation

We conducted sorting validation tests to ensure that device manufacturing and sorting was consistent between different devices and users. We used 1.53 μm polystyrene beads (Cat #17133, Polysciences Inc.) to mimic deformable red cells. The beads were suspended at 0.1% concentration in HBSS with 0.2% Pluronic F127 and 0.2% TWEEN-20 (MilliporeSigma) to prevent bead aggregation. The solution was input to the microfluidic device and run for 20 minutes. Images of the sorted beads were captured at the outlets and were manually counted using Fiji ImageJ. Users 1, 2, and 3 conducted 13 tests using devices from 5 different secondary master molds. We find intra- and inter-user microbead sorting was consistent (Fig. S1). Details are found in the Supplementary Materials.

### Image acquisition

Image scans of the blood smears were conducted using a 60× phase contrast objective and a DS-Qi2 camera on a Nikon Ti-2E inverted microscope using NIS Elements software. Illumination for the brightfield images was implemented using the built-in Ti-2E LED. Gain and exposure were automatically determined by NIS Elements for consistency and to avoid user bias. Vertical offset for focusing the images was determined using the built-in autofocus to create focus surfaces for each scan. These settings ensured consistent imaging parameters throughout the image acquisition. Components of the full image scan were 2424 × 2424 pixel BMP images with 14-bit depth.

### Segmentation and augmentation

The 2424 × 2424 pixel images were segmented into 81 non-overlapping images of 256 × 256 pixels. This produced 34,992 segmented images per class, which were cleaned to remove all images containing no cells. Prior to training, cell images were transformed including normalization, conversion to grayscale, and random horizonal flipping using Torchvision 0.11.1. Each class dataset was split 80:20 for the training and testing sets. From this split, 10,000 images were selected for training and 2,000 for testing. Further, 2,000 images from the training set were held-out for five-fold cross validation.

### Convolution neural network model

We designed a convolutional neural network (CNN) to conduct image feature extraction and classification using the PyTorch 1.10.0 library in Python 3.8.10. The network starts by accepting a 1-channel input of 256 × 256 pixels. The model begins with a 256-channel convolution layer with a kernel size of 7 × 7. The second layer is a 128-channel convolution layer with a kernel size of 5 × 5. The third convolutional layer has 64 channels, the fourth has 32 channels and the fifth has 16 channels, all with kernel sizes of 3 × 3. All convolutional layers are followed by ReLU activate, batch normalization, and max pooling with kernel size 2 × 2 and a stride of 2. Then, outputs of the convolutions are flattened into a 1-dimensional array for use in the fully connected layers. The model consists of 3 fully connected dense layers, each followed by ReLU activation, batch normalization, and 20% dropout. This procedure learns the identified features from the convolutional layers. For storage duration classification, the model outputs three nodes, one for each storage class (days 0, 7 and 14). Further, for deformability classification the model outputs four classes, one for each pre-determined deformability bin (RS < 2.50, RS 2.50-2.74, RS 2.75-2.99, or RS > 2.99). Alternatively, the regression model’s final layer is a linear layer with one output, the RS value prediction.

### Training environment

Our model was implemented for training, validation, and testing on the Compute Canada (Digital Research Alliance of Canada) Cedar cluster hosted at Simon Fraser University (Burnaby, Canada). This cluster consists of 94,528 CPU cores and 1,352 GPU devices. Most models were run with 32 GB of memory on 1 node with 8 tasks per node. In addition, training was conducted on a NVIDIA P100 Pascal GPU with 12G HBM2 memory. For the models trained with data from all donors, 2 to 3 CPU nodes were used. The training time was between ∼3 hours and ∼36 hours depending on the size of the datasets, number of epochs, learning rate, and other hyperparameters.

### Training

For each training session of the network, the data was passed through the model for 10 to 30 epochs with stochastic gradient descent (0.90 Nesterov momentum) and a learning rate between 0.001 and 0.01. The best combination of epochs and learning rate was found iteratively to find the best combination for training convergence and validation accuracy for each donor/dataset combination. Training ceased after there was no substantial loss improvements over the past 3 epochs. Batch sizes of 32 and 64 were used and the classification models utilized categorical cross entropy loss while the regression model used mean squared error loss. Further, the model was validated using five-fold cross validation. Reported training and validation accuracies are averaged over the five-folds.

### Testing

To further assess the models’ performances, testing occurred on separate datasets consisting of the 2,000 images held out per donor per class or label. In the collective donor storage duration classification case, the generalizability of the model was assessed by holding out each donor from training and using it for testing. Nine of these trials were conducted, one for each donor held-out.

## Ethics approval statement

This study was approved by the University of British Columbia’s Clinical Research Ethics Board (UBC REB# H19-01121) and Canadian Blood Services Research Ethics Board (CBS REB# 2019-029).

## Acknowledgments

We are grateful to Canadian Blood Services’ blood donors who made this research possible.

## Author contributions

H. M. supervised the study. H. M. and E.S.L. conceived the idea. E.S.L., Y.C., E.I., and K.M. performed the experimental work. E.S.L. performed the computational work. All authors wrote the manuscript.

